# Modeling and Predicting Cancer Clonal Evolution with Reinforcement Learning

**DOI:** 10.1101/2022.12.11.519917

**Authors:** Stefan Ivanovic, Mohammed El-Kebir

## Abstract

Cancer results from an evolutionary process that typically yields multiple clones with varying sets of mutations within the same tumor. Accurately modeling this process is key to understanding and predicting cancer evolution. Here, we introduce CloMu (<u>Clo</u>ne To <u>Mu</u>tation), a flexible and low-parameter tree-generative model of cancer evolution. CloMu uses a two-layer neural network trained via reinforcement learning to determine the probability of new mutations based on the existing mutations on a clone. CloMu supports several prediction tasks, including the determination of evolutionary trajectories, tree selection, causality and interchangeability between mutations, and mutation fitness. Importantly, previous methods support only some of these tasks, and many suffer from overfitting on datasets with a large number of mutations. Using simulations, we demonstrate that CloMu either matches or outperforms current methods on a wide variety of prediction tasks. In particular, for simulated data with interchangeable mutations, current methods are unable to uncover causal relationships as effectively as CloMu. On breast cancer and leukemia cohorts, we show that CloMu determines similarities and causal relationships between mutations as well as the fitness of mutations. We validate CloMu’s inferred mutation fitness values for the leukemia cohort by comparing them to clonal proportion data not used during training, showing high concordance. In summary, CloMu’s low-parameter model facilitates a wide range of prediction tasks regarding cancer evolution on increasingly available cohort-level datasets.

## 1 Introduction

Cancer results from an evolutionary process during which somatic mutations accumulate in a population of cells. This process results in a tumor composed of multiple sub-populations of cells, or *clones*, with varying sets of mutations [7]. As different clones within the same tumor harbor different sets of mutations, they have varying phenotypes and fitness [8]. Moreover, while each cancer results from a separate evolutionary process, there are repeated patterns of evolution that recur across cancers [9]. The key challenge in comparative cancer phylogenetics revolves around identifying such repeated evolutionary patterns or trajectories [10]. This is a challenging problem and requires a mathematical model that accurately captures the somatic evolutionary process of cancers.

There are many existing methods for modeling cancer evolution with single nucleotide variations, which can be distinguished into three classes (Table 1). The first class of methods are based on tree generative models, which include TreeMHN [1] and HINTRA [2]. The second class of methods are consensus tree models, which seek a small number of consensus trees that summarize common patterns among patient trees. Methods such as REVOLVER [5], RECAP [3] and CONETT [4] fall into this category. The third class of methods utilize statistical tests to evaluate patterns of co-occurrence and mutual exclusivity of mutations across trees without trying to fully model the evolutionary process. GeneAccord [6] is one such method. While consensus tree methods have the advantage of being able to detect complicated patterns in tumor evolution, tree generative models must be carefully designed to accommodate complex multi-mutation effects. An example of such an effect is when mutations *s* and *t* only increase the probability of mutation *r* when they are present together. Conversely, tree-generative methods have the advantage of being able to directly model how a clone’s mutations affect the probability of new mutations occurring.

**Table 1:** Overview of methods for detecting patterns of cancer evolution. For each method, we specify, from left to right, the underlying tree model, the number of parameters, support for evolutionary trajectories, tree selection, multi-mutation effects, signed and bidirectional causality inference, mutation fitness, interchangeable mutations and their pathways. ^*^GeneAccord does not contain a generative model of trees, nor does it seek to infer a tree. Rather it takes as input a set of trees, and assesses the causality between all pairs of mutations using a statistical test. ^†^Indicates that while the feature is unimplemented, it could be supported by TreeMHN’s model.

| Method | Tree model | #Params | Evo. traj. | Tree selection | Multi-mut. effects | Signed causality | Bidir. causality | Mut. fitness | Int. muts |
| --- | --- | --- | --- | --- | --- | --- | --- | --- | --- |
| CloMu | generative | $O(mL)$ | ✓ | ✓ | ✓ | ✓ | ✓ | ✓ | ✓ |
| TreeMHN [1] | generative | $O(m^2)$ | ✓ | ✓/✗<sup>†</sup> | ✗ | ✓ | ✓ | ✓/✗<sup>†</sup> | ✗ |
| HINTRA [2] | generative | $O(m2^m)$ | ✓ | ✓ | ✓ | ✗ | ✓ | ✗ | ✗ |
| RECAP [3] | consensus | $O(m^2)$ | ✓ | ✓ | ✓ | ✗ | ✓ | ✗ | ✗ |
| CONETT [4] | consensus | $O(m^2)$ | ✓ | ✓ | ✓ | ✗ | ✓ | ✗ | ✗ |
| REVOLVER [5] | consensus | $O(m^2)$ | ✓ | ✓ | ✓ | ✗ | ✓ | ✗ | ✗ |
| GeneAccord [6] | statistical test* | $O(m^2)$ | ✗ | ✗ | ✗ | ✓ | ✗ | ✗ | ✗ |

A second distinguishing feature of current methods is how the number of model parameters scales with increasing number *m* of mutations (Table 1). Having the number of parameters scale too rapidly with the number of mutations can lead to greatly overfitting on real datasets as well as a prohibitive running time. One such example is HINTRA [2], where the number of parameters grows exponentially in *m*, and, consequently, it can only accommodate a handful of mutations. On the other hand, models that directly measure relationships between pairs of mutations or fit trees with edges determined by pairs of mutations have their parameters grow quadratically in the number of mutations [1, 3–6].

Finally, current methods can be distinguished by the prediction tasks they support. The majority of current methods, spanning both tree-generative [1, 2] and consensus tree methods [3–5], aim to identify evolutionary trajectories, which correspond to ordered sequences or trees of mutations reflecting repeated patterns of evolution. Additionally, several methods [2–5] use signals from across patients to resolve uncertainty in phylogeny inference within a single patient. Another task is the prediction of causality between pairs of mutations. This can be done in signed manner, distinguishing between causal and inhibitory relationship, as well as in a bidirectional manner, distinguishing between a mutation *s* causing *t* and vice versa. While TreeMHN [1] supports both signed and bidirectional causal relationships, GeneAccord [6] only supports signed causal relationships. On the other hand, consensus tree methods [3–5] are unable to support inhibitory causal relationships.

Here, we identify three additional tasks that no current evolutionary trajectory method supports. First, determining the fitness of a mutation, i.e. assessing how having that mutation impacts the rate at which a clone develops. Second, determining sets of interchangeable mutations that have identical impacts on subsequent mutation evolution. Third, determining pathways of interchangeable mutations that describe a collection of evolutionary trajectories between sets of interchangeable mutations. To accommodate these new and all previous tasks, we introduce CloMu (Clone To Mutation, Fig 1). Underlying CloMu is a tree-generative model which uses a neural network trained using reinforcement learning. Specifically, CloMu utilizes a low parameter two layer neural network with *L m* hidden neurons. This results in less parameters, i.e. *O* (*L* « *m*), than all current models, while maintaining a high flexibility to model complex multi-mutation effects (Table 1). Using simulations with known ground truth, we demonstrate that CloMu outperforms existing methods on several prediction tasks. Additionally, we apply CloMu to analyze a breast cancer [11] and acute myeloid leukemia cohort [12], identifying variability in mutation fitness as well as uncovering causal relationships and interchangeable mutations.

**Figure 1.**
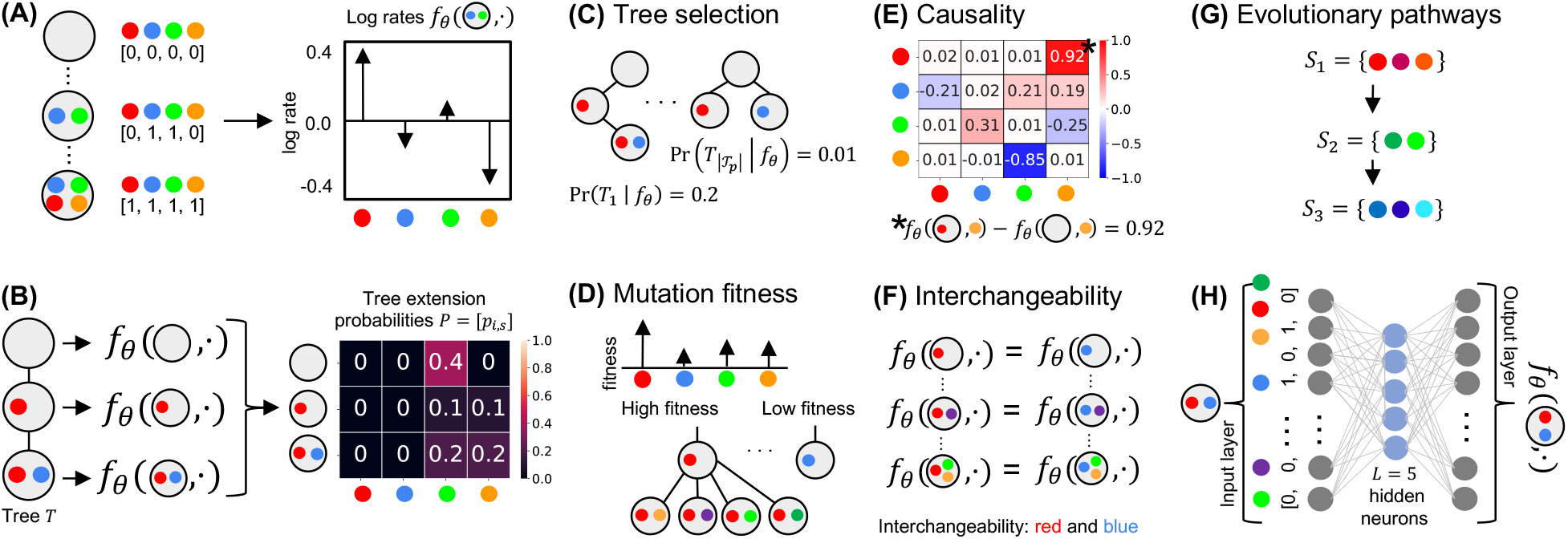
Overview of CloMu. (A) Using the independent clonal evolution assumption, our model determines a log rate *f*_*θ*_(**c**, *s*) of any clone **c** ∈ {0, 1 }^*m*^ acquiring a mutation *s* ∈ [*m*]. (B) This in turn enables us to compute probabilities *P* = [*p*_*i,s*_] that the next mutation to occur on a tree *T* is *s* at the node/clone **c**_*i*_. The resulting Independent Clonal Evolution problem seeks model parameters *θ* that maximize the data probability Pr(𝒯 _1_, …, 𝒯_*n*_ |*f*_*θ*_) of a cohort of trees for *n* patients. (C-G) We use the model for five prediction tasks: (C) tree selection for each patient, (D) determination of mutation fitness, (E) causality inference for pairs of mutations, (F) identifying interchangeable mutations and (G) their evolutionary pathways. (H) CloMu represents *f*_*θ*_ using a low-parameter, two-layer neural network that is trained via reinforcement learning, achieving flexibility to capture complex interactions and multimutation effects while avoiding overfitting on datasets with a large number of mutations.

## 2 Problem Statement

### Input and output

We take as input tumor phylogenies of *n* patients with *m* total mutations. Due to uncertainty in tree inference from sequencing data [13], each patient *p* has a set _*p*_ of multiple possible trees 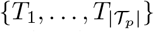. Similarly to HINTRA [2] and TreeMHN [1], we make the *independent clonal evolution* assumption that the event of a clone acquiring a new mutation only depends on the genotype of that clone. While clonal cooperation has been described in the literature [14, 15], the resulting problem under our assumption still enables meaningful analyses even for cases with violations to this assumption (Appendix B.6). Each clone can be represented as a binary vector **c** ∈ {0, 1}^*m*^. The goal is to identify a model *f*_*θ*_ that best describes the causal relationship between clones **c** and acquired mutations [*m*] = {1, …, *m*} under our assumption for the observed data {𝒯_1_, …, 𝒯 }_*n*_.

### Generative model

Our generative model for trees is similar to the generative models employed by HINTRA [2] and TreeMHN [1], with a couple of differences that we will discuss below. Per our assumption, the probability of a new mutation *s* ∈ [*m*] occurring on any clone **c** ∈ {0, 1 }^*m*^ is only a function of **c** and *s*. Specifically, we model this using the function *f*_*θ*_ : {0, 1 }^*m*^ × [*m*] →ℝ, which outputs the logarithm of the rate at which the mutation occurs on the clone (Fig. 1A). As such, the probability of mutation *s* occurring on the clone **c** in the next *δ* time units is *δ* exp(*f*_*θ*_(**c**, *s*)) for small *δ*. Starting with an initial tree *T* ^(0)^ with just a single node corresponding to the normal clone **c**_0_ = [0, …, 0]^⊤^, we sample the mutation that occurs next, yielding a new tree *T* ^(1)^ with an additional clone **c**_1_ that introduces one mutation w.r.t. **c**_0_. More precisely, given a tree *T* ^(*k*)^ with clones *C*^(*k*)^ = [**c**_0_, …, **c**_*k*_]^⊤^, the conditional probability Pr((**c**_*i*_, *s*) | **c**_0_, …, **c**_*k*_, *f*_*θ*_) of the next mutation *s* to occur on clone **c**_*i*_ among clones *C*^(*k*)^, denoted by (**c**_*i*_, *s*), equals

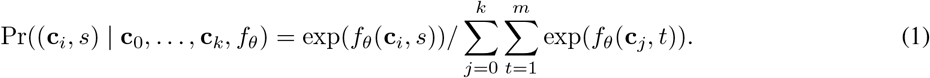

This yields a new tree *T* ^(*k*+1)^ with a new clone **c**_*k*+1_ with **c**_*i*_ as its parent. Equivalently, we may represent the conditional probability density function with a softmax as *P* = [*p*_0,1_, …, *p*_*k,m*_] ⊤ = softmax([*f*_*θ*_(**c**_0_, 1), …, *f*_*θ*_(**c**_*k*_, *m*)]^⊤^) where *p*_*i,s*_ = Pr((**c**_*i*_, *s*) | **c**_0_, …, **c**_*k*_, *f*_*θ*_) — see Fig. 1B. Optionally, the infinite sites assumption can be enforced by setting the probabilities *p*_0,*s*_, …, *p*_*k,s*_ of a new mutation *s* to zero if mutation *s* already exists in one of the clones **c**_0_, …, **c**_*k*_. We terminate the tree generation process when we reach a pre-specified number, 𝓁 ≤ *m* of mutations.

### Parameter estimation

Our goal is to find the model parameters *θ*^*^ that maximize the probability of observing the input data, i.e., *θ*^*^ = argmax_*θ*_ Pr(𝒯_1_, …, 𝒯_*n*_ |*θ*). To do so, we must estimate the probability of any tree *T*. However, there are multiple possible ways of generating each tree. For example, consider the tree *T* with 𝓁 = 2 mutations having the normal clone **c**_0_ as the root with two children: a clone with mutation 1 and a clone with mutation 2. There are two ways to generate *T*, either mutation 1 or mutation 2 could occur first on **c**_0_ followed by the other mutation, i.e., (**c**_0_, 1), (**c**_0_, 2) or (**c**_0_, 2), (**c**_0_, 1). We refer to a specific way of generating a tree *T* with 𝓁 mutations as a *tree generating process G*, which is an ordered list 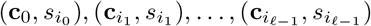 where each pair indicates the source clone and the mutation that was added to it (Fig. S1). Using the independent clonal evolution assumption, we can now express the probability of a tree generating process 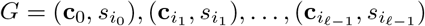 as

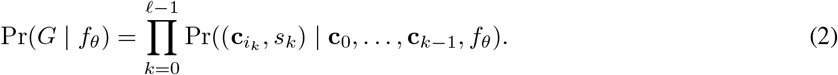

We denote the set of all tree generating processes that yield a tree *T* by 𝒢 (*T*) such that Pr(*T* | *f*_*θ*_) = ∑_*G*∈𝒢 (*T*)_ Pr(*G* | *f*_*θ*_). As tumors are separate evolutionary processes, we have independence across patients and can therefore express the probability of our data as

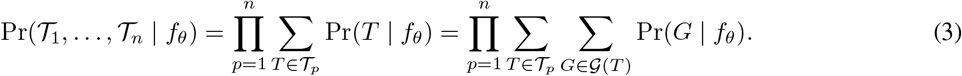

This leads to the following problem.

**Problem 1** (Independent Clonal Evolution). Given a cohort of tumor phylogenies 𝒯_1_, …, 𝒯_*n*_ for *n* tumors on *m* mutations and model class *f*, find model parameters *θ* such that Pr(𝒯_1_, …, 𝒯_*n*_ | *f*_*θ*_) is maximum.

We note that HINTRA [2] and TreeMHN [1] solve the above problem with different model classes *f*. The function *f*_*θ*_ employed by HINTRA explicitly enumerates all 2^*m*^ × *m* clone-mutation pairs. The advantage of HINTRA’s model is its flexibility to fit any pattern of clone-to-mutation interactions; but this comes at the expense of severely restricting its scalability to a small number of mutations. On the other hand, TreeMHN defines *f*_*θ*_ using a linear model of additive rate coefficients for all *m* × *m* pairs of mutations. As we will show later in Section 4.1, this scales well but does not capture non-additive clone-to-mutation interactions, nor shared properties of mutations such as two mutations similarly affecting the probability of other mutations. As discussed in Section 3.2, in our method CloMu, the model *f*_*θ*_ is a two-layer neural network with *L* « *m* hidden neurons. The smaller number of parameters and the ability to model non-additive clone-to-mutation interactions leads CloMu to achieve flexibility like HINTRA while also being more resistant to overfitting than either TreeMHN or HINTRA on datasets with a large number of mutations (Table 1). Beyond the model class *f*, there are other subtle differences. Rather than considering the set 𝒢 (*T*) of all tree generative processes of a tree *T*, HINTRA considers a single tree generative process *G* to compute the probability of *T*. Moreover, in TreeMHN, the number, 𝓁 of mutations in a patient’s tumor is determined by the model rather than taken as input.

## 3 Method

### 3.1 Prediction Tasks

The model described in Section 2 can be used in several ways to gain insights into cancer evolution. First, one can use the model to reduce uncertainty in tree inference by determining which phylogenies are most likely to be the true phylogeny for a particular patient. This would be done by selecting trees *T* ∈ 𝒯_*p*_ with a high value of Pr(*T* | *f*_*θ*_) for each patient *p* (Fig. 1C).

Second, the model can be used to determine the fitness of different mutations based on how they effect the likelihood of new mutations occurring on a clone (Fig. 1D). We do so by again considering the set {**c**_1_, …, **c**_*m*_ } of clones where each clone **c**_*s*_ consists of only mutation *s*. Intuitively, the fitness of a clone **c**_*s*_, and consequently mutation *s*, is proportional to the rate at which a mutation would occur on the clone. As such, normalizing this over all clones **c**_1_, …, **c**_*m*_ yields the expression of *mutation fitness*:

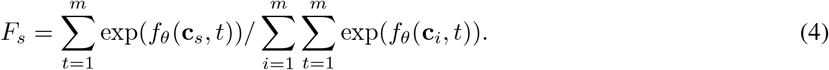

Third, the model can be used to infer causal relationships (Fig. 1E). We will define absolute and relative causality by considering the set {**c**_1_, …, **c**_*m*_} of clones where each clone **c**_*s*_ contains only mutation *s*. Recall that *f*_*θ*_(**c**, *s*) is the logarithm of the rate at which a mutation *s* occurs on clone **c**. For absolute causality, we have when *f*_*θ*_(**c**_*s*_, *t*) −*f*_*θ*_(**c**_0_, *t*) is positive, the mutation *s* increases the rate of mutation *t* relative to the normal clone. Conversely, when *f*_*θ*_(**c**_*s*_, *t*) − *f*_*θ*_(**c**_0_, *t*) is negative, the mutation *s* decreases the rate of mutation *t* relative to the normal clone. If *f*_*θ*_(**c**_*s*_, *t*) − *f*_*θ*_(**c**_0_, *t*) = 0, there is no causal relationship between from *s* to *t*. In practice, some threshold value *τ* > 0 in the strength of the causal relationship must be used in order to avoid false positives. Therefore, for absolute causality, we say *s causes t* if *f*_*θ*_(**c**_*s*_, *t*) − *f*_*θ*_(**c**_0_, *t*) > *τ, s inhibits t* if *f*_*θ*_(**c**_*s*_, *t*) − *f*_*θ*_(**c**_0_, *t*) < −*τ*, and there is no causal relationship from *s* to *t* otherwise. Note that causality is bidirectional, i.e., we can assess causality from *t* to *s* by evaluating *f*_*θ*_(**c**_*t*_, *s*) −*f*_*θ*_(**c**_0_, *s*), amounting to a total of 9 possible pairs of causal relationships between any two mutations. A drawback of absolute causality is that a mutation *s* is highly fit (large *F*_*s*_), it may have a positive causal relationship with all other mutations according to our definition of absolute causality (Table S1). To account for this, we define *relative causality* from *s* to *t* in terms of

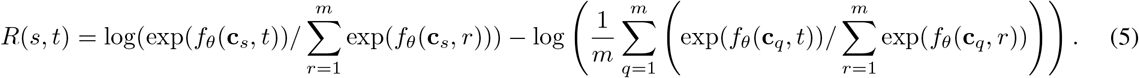

This measures how much the mutation *t* increases the relative probability that *s* will be the next mutation to occur on the clone, relative to the normal clone **c**_0_. Similarly to absolute causality, we say *s causes t* if *R*(*s, t*) > *τ, s inhibits t* if *R*(*s, t*) < −*τ*, and there is no causal relationship from *s* to *t* otherwise.

Fourth, the model can be used to detect shared properties between mutations, including interchangeable mutations (Fig. 1F). Two mutations *s* ≠ *t* are *interchangeable* if it holds that *f*_*θ*_(**c**, *r*) ≈ *f*_*θ*_(**c**′, *r*) for all mutations *r* and for any two clones **c** and **c**′ that are identical except for **c** containing mutation *s* but not *t*, and vice versa for **c**′. One shared property of interest is when two mutations incur the same set of causal relationships with other mutations but with different effect strengths, i.e., *f*_*θ*_(**c**, *r*) ≈*α* · *f*_*θ*_(c′, *r*) for some *α* > 0 and all mutations *r* ∈ [*m*]. Another property is when two mutations have the same causal relationships on some subset of mutations, i.e., *f*_*θ*_(**c**, *r*) ≈ *f*_*θ*_(**c**′, *r*) where *r* ∈ *R* ⊆ [*m*].

Fifth, we can determine pathways of interchangeable mutations (Fig. 1G). Specifically, the model is capable of determining pathways in which mutation *s* causes mutation *t*, and mutation *s* and *t* are required together in order to cause mutation *r*. Mathematically, the latter would result in *f*_*θ*_(**c**_*s*_, *r*) ≈ *f*_*θ*_(**c**_0_, *r*) ≈ *f*_*θ*_(**c**_*t*_, *r*) and *f*_*θ*_(**c**_*st*_, *r*) > *f*_*θ*_(**c**_0_, *r*) where **c**_*st*_ is the clone that contains only mutations *s* and *t*. Additionally, the model generalizes to the case where some set *A* of mutations causes all mutations from another set *B*, and pairs of mutations in the two sets are required together in order to cause mutations *C*.

### 3.2 CloMu: Low-parameter Neural Network Trained via Reinforcement Learning

We introduce CloMu (<u>Clo</u>ne To <u>Mu</u>tation), which uses a two layer neural network with a small number *L* of hidden neurons for the function *f*_*θ*_ (Fig. 1H). This model has 2*Lm* + *m* + *L* = *O*(*Lm*) total parameters. In experiments in this paper, we use *L* = 5 hidden neurons unless stated otherwise. As the number of parameters grows linearly in the number of mutations, this model is very resistant to overfitting. Moreover, we made a slight modification to the neural network to allow for easier regularization, which is used primarily for the sake of interpretability but also used to avoid overfitting on some extremely small datasets. This modification did not affect the scaling of the number parameters. We refer to Appendix A.1 for more details.

The small number *L* of hidden neurons allows us to form a small *L*-dimensional representation of any input clone **c**. Applying this to the set of clones with only one mutation, then allows us to generate small latent representations of mutations, which can be used for determining mutation interchangeability. Specifically, the latent representation encapsulates any effect the mutation could have on some clone according to the model, and thus measuring the similarity of two latent representations also measures the similarity between the two mutations. In practice, we use the Euclidean distance between latent representations to determine which mutations are interchangeable. Note, the median value across all mutations is subtracted from each component of each latent representation for the sake of interpretability.

For the task of modeling evolutionary pathways of interchangeable mutations, it is vital to understand when two mutations are required together in order to cause a third mutation. Having a non-linear model such as our neural network is thus required for this task. Specifically we find evolutionary pathways of interchangeable mutations through the following procedure described extensively in Appendix A.2. First, we evaluate the probability of the tumor evolving in some evolutionary pathway according to our model. Then, we evaluate the probability of the tumor evolving in that same pathway under a null hypothesis where mutations have no effect on clonal evolution. Finally, we search for evolutionary pathways where the probability according to our model vastly exceeds the null hypothesis probability.

The sets 𝒯_1_, …, 𝒯_*n*_ and especially 𝒢 (*T*) can be very large. For instance, if *T* is the star tree where 10 mutations are made on the root node, 𝒢 (*T*) will have 10! = 3,628,800 elements. Explicitly calculating *P* (*G* |*θ*) for every *G* ∈ 𝒢 (*T*) and *T* ∈ 𝒯_*p*_ can be infeasible. To avoid doing this, we use reinforcement learning to train the neural network, i.e., infer model parameters *θ* that maximize the likelihood of the data. To accomplish this, we use policy learning where the model is given a reward when it generates a tree in the dataset. Specifically, the reward is proportional to how much increasing the probability of that tree would increase the overall log probability of our dataset. Typically in reinforcement learning, the reward given to the model for a given action sequence is independent of the model itself. In our case, in order to maximize the probability of observing the data, our reward is a function which is dependent on the model. However, it is critical to correctly optimizing our objective that the rewards are treated only as numerical values and that the gradient of the reward is not taken with respect to the parameters. As is typical, our model is trained via stochastic gradient descent on the typical policy learning loss function. Additionally, we have a modified our sampling procedure to increase training speed and accuracy in the case that there are few trees per patient, and a consequently adjusted reward function. We refer to Appendix A.3 for more details.

We implemented CloMu in Python 3 using the PyTorch library. CloMu is open source and available at https://github.com/elkebir-group/CloMu.

## 4 Results

### 4.1 Simulations

We use simulations with known ground truth to assess the performance of CloMu and several existing methods on the prediction tasks we outlined in Section 3.1. We consider four sets of simulation instances: (i) simulations to assess tree selection and causality, (ii) simulations to assess mutation interchangeability and evolutionary pathways as well as previous simulations generated in the (iii) RECAP [3] and (iv) TreeMHN [1] papers (Table 2). We include TreeMHN [1], RECAP [3], REVOLVER [5] and GeneAccord [6] in the benchmarking, and refer to Appendix B.2 for parameter settings.

**Table 2:** Characteristics of simulation datasets. We considered six sets of simulations, labeled (I-a) to (IV) with varying number of driver mutations, total number of mutations, number of interchangeable mutation sets, number of interchangeable mutations per set, number of patients, the presence of both causation and inhibition, the number of evolutionary pathways of interchangeable mutations and the total number of instances. Simulations (III) correspond to previously published RECAP simulations [3]. Simulations (IV) correspond to previously published TreeMHN simulations [1].

|  | # Drivers | # Mutations | # Int. mut. sets | # Int. muts per set | # Patients | Signed causality | # Pathways | # Instances |
| --- | --- | --- | --- | --- | --- | --- | --- | --- |
| (I-a) | 5 | 10 | NA | NA | 500 | no | NA | 20 |
| (I-b) | 10 | 10 | 5 | 2 | 400 | yes | NA | 20 |
| (I-c) | 15 | 15 | 5 | 3 | 600 | yes | NA | 20 |
| (II) | $\{3, \dots, 18\}$ | 20 | # Pathways $\times 3$ | $\{1, 2, 3\}$ | 500 | no | $\{1, 2\}$ | 30 |
| (III) | NA | $\{5, 12\}$ | NA | NA | $\{50, 100\}$ | NA | NA | 400 |
| (IV) | $\{10, 15, 20\}$ | $\{10, 15, 20\}$ | NA | NA | 300 | yes | NA | 60 |

We begin by discussing the generation of the first set of simulation instances to assess tree selection, causality and mutation interchangeability. We focus on the simplest case with no interchangeable mutations. We considered *m* = 10 mutations, subdivided into five driver mutations and five passenger mutations. For every ordered pair (*s, t*) of distinct driver mutations, there is a 50% chance that *s* causes *t*. Let *X* be the resulting set of causal relationship pairs. Note that |*X*| ≤ 20 as there are 20 ordered pairs (*s, t*) of distinct driver mutations. For each pair (*s, t*) ∈ [*m*] × [*m*] of mutations, we set the rate multiplier *g*(*s, t*) = 11 if (*s, t*) ∈ *X* and *g*(*s, t*) = 1 otherwise. We defined the rate *f* (**c**, *t*) of a clone **c** ∈ {0, 1}^*m*^ acquiring a mutation *t* as 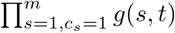. Given a number *n* = 500 of patients, we generated one ground-truth tree 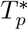 for each patient *p* ∈ [*n*] by first drawing the number, 𝓁 of mutations of tree 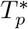 uniformly from {5, 6, 7}. We used the rates *f* (**c**, *t*) to construct 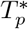 following the generative process with, 𝓁 mutations discussed in Section 2 — with exp(*f*_*θ*_(**c**, *t*)) replaced with *f* (**c**, *t*). Although not required by CloMu, we imposed the infinite sites assumption in our simulations. Finally, we simulated five bulk DNA sequencing samples and performed clonal tree enumeration using SPRUCE [16], resulting in sets 𝒯_1_, …, 𝒯_*n*_ of trees per patient with varying number of trees per patient (Fig. S2A). We generated a total of 20 simulation instances, denoted Simulations (I-a). Additionally, to assess the effect of interchangeable mutations, we generated two additional sets of 20 simulation instances with 5 pairwise disjoint sets of 2 and 3 interchangeable mutations, denoted Simulations (I-b) and (I-c), respectively. Finally, we generated of a set of 20 simulation instances, denoted Simulation (II), to assess evolutionary pathway identification. We refer to Table 2 and Appendix B.1.1 for further details.

For the tree selection task, we compared CloMu to RECAP and REVOLVER on Simulations (I-a) (Table 2). For each simulation instance, we determined the tree selection accuracy defined as the fraction of patients for which each method correctly identified the ground-truth tree. Fig. 2A shows that CloMu achieves the highest tree selection accuracy (median: 0.76), followed by REVOLVER (median: 0.70) and then RECAP (median: 0.65). To assess causality inference, we compare against TreeMHN and GeneAccord, which directly support causality inference (Table 1). Although RECAP and REVOLVER do not directly support this task, we used a simple heuristic based on the frequency of mutation pairs/edges (*s, t*) among selected trees (Appendix B.3). Fig. 2B shows that CloMu and TreeMHN performed near perfectly on this task (median recall and precision of 1.0), followed by RECAP and REVOLVER (median recall: 0.76 and 0.77, and median precision: 1.0 and 1.0, respectively) and then GeneAccord (median recall: 0.78 and median precision: 0.85).

**Figure 2.**
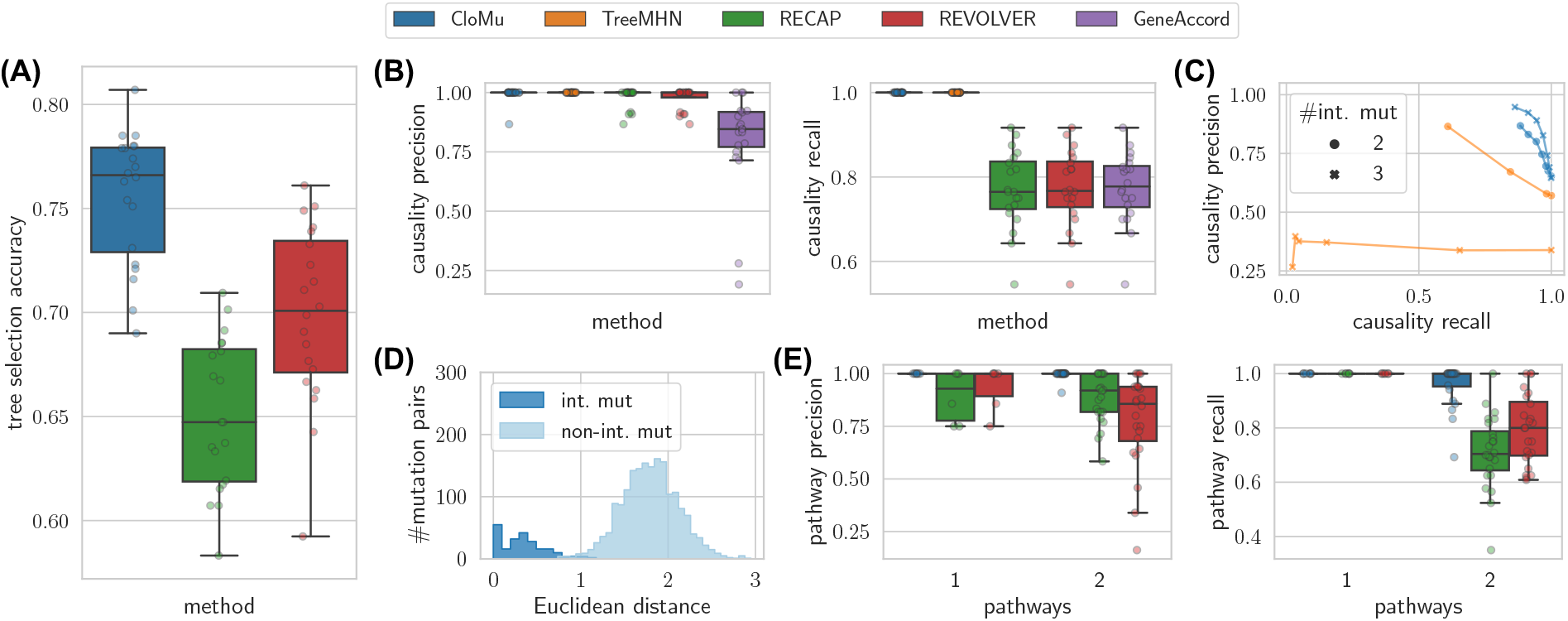
CloMu outperforms existing methods on several prediction tasks on simulated data with known ground truth. (A) Tree selection accuracy measures the ability to correctly identify the ground-truth tree from a set 𝒯_*p*_ of possible trees generated from simulated bulk sequencing for each patient *p*. (B) Causality precision and recall measure the ability to determine positive causal relationships between ordered pairs of mutations. Panels (A) and (B) shows results for Simulations (I-a). (C) On Simulations (I-b) and (I-c), causality precision and recall measure the ability to identify to identity causation and inhibition between pairs of mutations in the presence of mutation interchangeability. (D) On Simulations (II), interchangeability detection demonstrates that the latent representations generated by our model are meaningful, accurately encapsulating mutation similarity. (E) On simulations (II), pathway precision and recall measure the ability to determine evolutionary pathways in the presence of both interchangeability and complex multi-mutation interactions.

Upon observing CloMu and TreeMHN’s near perfect performance, we decided to include interchangeable mutations as well as inhibitory relationships (Simulations (I-b) and (I-c), see Table 2). We refer to Appendix B.1.2 for the updated definitions of causality precision and recall for the case of signed causality, which match TreeMHN’s definitions. We found that CloMu maintained good performance whereas TreeMHN’s performance dropped, especially with increasing numbers of interchangeable mutations (Fig. 2C). We believe the reason TreeMHN performed poorly on these instances is its assumption of independent causal relationships between pairs of mutations. The number *m*^2^ − *m* of causal relationships TreeMHN must independently determine grows quadratically with the number *m* of mutations. In contrast, if CLoMu’s neural network learns that there are only *k* « *m* non-interchangeable types of mutations, it must only detect *k*^2^ causal relationships. Specifically, for these simulation instances there were *k* = 5 sets of interchangeable mutations among a total of *m* ∈{10, 15 } mutations (see Appendix B.1.1), leading to *k*^2^ = 25 causal relationships for CloMu to detect and *m*^2^ −*m* ∈ {90, 210 }for TreeMHN.

We now focus on Simulations (II) to further assess performance on determining interchangeable mutations and their evolutionary pathways. Fig. 2D shows that CloMu accurately distinguishes between pairs of interchangeable and non-interchangeable mutations as assessed by the Euclidean distance on CloMu’s latent representation ℝ^*L*^ of *L* = 5 hidden neurons. We note that these simulations contain causal effects that only occur in the presence of specific combinations of mutations on a clone (Appendix B.1.1). Accurately determining the pathways requires modeling these multi-mutation effects. In order to have some baseline for comparison, we applied a heuristic to determine pathways of interchangeable mutations from the trees selected by RECAP and REVOLVER (Appendix B.3). Briefly, our heuristic measured the number of times each edge occurs in selected trees for this method, and then inferred the pathway edges as all edges which occur above some frequency. TreeMHN and GeneAccord could not be adapted to form a baseline because they do not select trees. Additionally, they have no way of modeling effects which only occur in the presence of a combination of mutations. We determined the number of true and false positives as well as false negatives by comparing the set of true pathways edges and predicted edges, enabling us to compute pathway precision and recall. We found that CloMu, RECAP, and REVOLVER all achieved a median pathway recall of 1.0 in simulations with only one pathway (Fig. 2E). However, CloMu outperformed the baselines in terms of precision (overall median: 1.0 vs. 0.92 and 0.87 for RECAP and REVOLVER, respectively), and especially for cases with two evolutionary pathways, it additionally outperformed the baseline methods in terms of recall (overall median: 1.0 vs. 0.75 and 0.83 for RECAP and REVOLVER, respectively).

Finally, we ran CloMu on data generated in the RECAP and TreeMHN papers, denoted as Simulations (III) and (IV), respectively (Table 2). We found that we matched RECAP’s performance (Appendix B.4) but that CloMu was outperformed by TreeMHN when using *L* = 5 hidden neurons but achieved approximately similar performance when using a linear model with no hidden neurons (Appendix B.5). It is important to note that that this results from the absence of interchangeable mutations or any mutations with shared causal properties in TreeMHN’s simulations. Such mutations are present in real data as we will show. As previously stated, our model utilizes the independent clonal evolution assumption. To demonstrate that our model is robust to violations in this assumption, we generated a causal relationship simulations with clonal cooperation, demonstrating robustness by maintaining high causal inference performance on these data (Appendix B.6).

In summary, this simulation study demonstrates that CloMu is a versatile model and supports a wide variety of prediction tasks. CloMu’s low-parameter neural network affords it the flexibility to model complex multi-mutation interactions while retaining resistance to overfitting.

### 4.2 Breast Cancer Cohort

We applied CloMu to a breast cancer cohort composed of 1918 tumors from 1756 patients that were sequenced using a gene panel. We used the same processing steps as in the RECAP paper [3], restricting our analysis to singlenucleotide variants that occur in copy neutral autosomal regions followed by running SPRUCE [16] to obtain a set 𝒯_*p*_ for each patient *p*. As in the RECAP analysis, we only retained mutations that occurred in at least 100 patients and removed patients that did not contain any of these recurrent mutations. In order to use the RL algorithm that considers all trees rather sampling them (Appendix A.3.3), we additionally removed patients with more than 9 mutations and consequently had a large number of trees. This left us with a dataset with *n* = 1224 patients and *m* = 406 mutations. We ran CloMu with defaults settings and *L* = 5 hidden neurons. Our RL algorithm took less than three hours to train the neural network on a laptop with a 2.4 GHz CPU and 64 GB of RAM without using a GPU.

On the task of determining mutation fitness, we found that only eight mutations have high fitness values (*>* 0.003) as shown in Fig. 3. Specifically, the highest fitness mutation was determined to be *TP53*, a known tumor suppressor gene [17]. The clone with only *TP53* had a fitness (0.0125) over 5 times as the median fitness value (0.00234). Additionally, in order, the next highest fitness mutations were determined to be *CDH1, PIK3CA, GATA3*, and *MAP3K1*, with fitness values between 0.00658 and 0.0104. Finally, the third tier consists of *ARID1A, ESR1* and *KMT2C* with fitness values between 0.00384 and 0.00440. All of these are known driver mutations [18].

**Figure 3.**
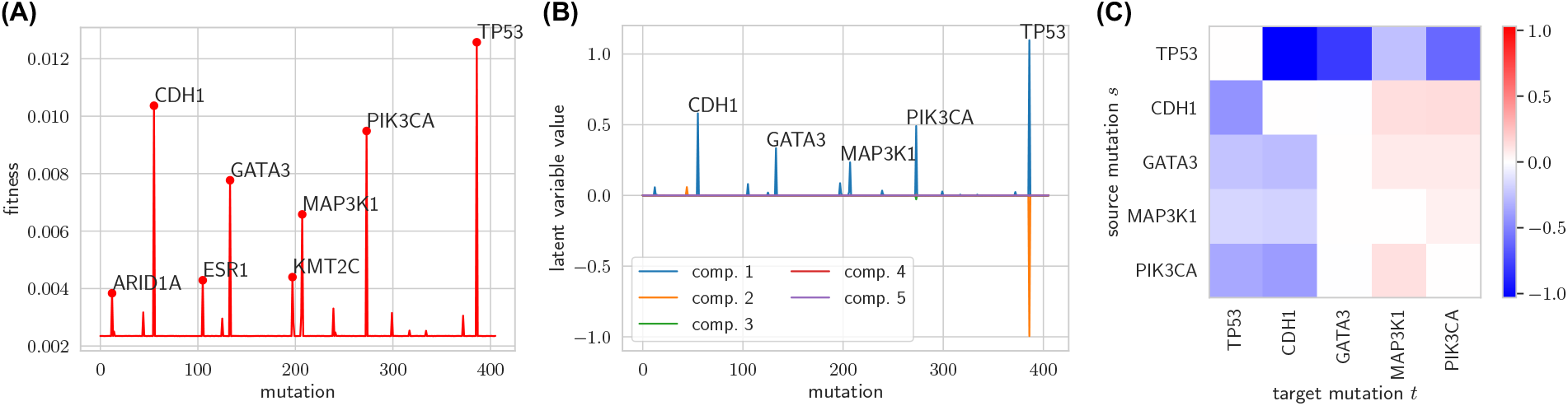
CloMu determines mutation fitness, uncovered mutation similarity and interchangeability, and finds relative causal relationships on a breast cancer cohort [11]. (A) CloMu identified eight mutations which have a far greater fitness value than other mutations. (B) Among these eight mutations, five mutations had latent representations with significant nonzero magnitudes. This plot shows that *CDH1* and *PIK3CA* as well as *GATA3* and *MAP3K1* are interchangeable on these data. (C) Relative causal relationships between mutations pair (*s, t*). A value greater than 0 (red) is indicative of mutation *s* (row) causing mutation *t* (column) in the same clone, whereas a value smaller than 0 (blue) is indicative of mutation *s* inhibiting the occurrence of mutation *t* in the same clone.

On the task of determining interchangeability and shared mutation properties, we inspected the latent representations of all mutations. We found five mutations with significant magnitude latent representations, corresponding to the five highest fitness mutations: *CDH1, GATA3, MAP3K1, PIK3CA* and *TP53*. Among these mutations, we found two pairs of interchangeable mutations (Fig. 3B). First, *CDH1* and *PIK3CA* had similar latent representations, with similar values (0.580 and 0.491, respectively) in the first component and values of roughly 0 (magnitude under 0.03) in the other components. Second, *GATA3* and *MAP3K1* have similar latent representations, with similar values (0.334 and 0.233, respectively) in the first component and values of roughly 0 (magnitude under 0.001) in the other components. In fact, the only high-fitness mutation that our model determined to be highly unique is *TP53*. We found *TP53* to have a separate property expressed by having a different dimension in its latent representation unseen by other mutations (corresponding to the second component).

Finally, we analyzed relative causality among the five highest-fitness mutations. Recall that a relative causality value *R*(*s, t*) significantly greater than 0 is indicative of mutation *s* causing mutation *t* in the same clone whereas a value significantly smaller than 0 is indicative of mutation *s* inhibiting the occurrence of mutation *t* in the same clone (Section 3.1). We identified more negative (11 pairs) than positive (6 pairs) causal relationships between pairs (*s, t*) of mutations (Fig. 3C). This indicates that high-fitness mutations, or drivers, increase the likelihood of passenger mutations by more than they increase the likelihood of other driver mutations. This is especially the case for *TP53*, which has a median relative causality value of − 0.253 for the seven other highest-fitness mutations vs. a relative causality value of 0.476 for the remaining mutations. Among the positive causal relationships, we found that *CDH1* and *GATA3* both cause *PIK3CA*. This is in agreement with TreeMHN’s conclusions that *CDH1* causes *PIK3CA*, and that there is some level of co-occurrence between *GATA3* and *PIK3CA* [1].

### 4.3 Acute Myeloid Leukemia Cohort

We analyzed a cohort of 123 acute myeloid leukemia (AML) patients that underwent high-throughput single-cell DNA panel sequencing [12]. Due to the relatively few number of patients, we only analyzed the gene-level data with the exception of the gene *FLT3*, where we additionally distinguished an internal tandem duplication mutation in *FLT3* denoted as *FLT3-ITD*. We used the phylogenies inferred by Morita et al. [12] using SCITE [19]. We also restricted our analysis to the 77 patients with reported clonal prevalences. Again for training efficiency reasons for our particular RL implementation (Appendix A.3.3), we removed patients with more than 10 mutated genes. We arrived at *n* = 75 patients with *m* = 22 total mutated genes. One reason we chose to analyze this dataset is that it includes clonal prevalence data, an independent source of fitness information which we used for orthogonal validation of our fitness predictions. Because we collapsed mutations to the gene level, we disabled the infinite sites mode in CloMu. We used default parameters with *L* = 5 hidden neurons. Training the neural network took less than 1 hour on a laptop with a 2.4 GHz CPU and 64 GB of RAM without using a GPU.

We used CloMu to predict mutation fitness, interchangeability, and causality. Our model determined the most fit mutations to be *NPM1* and *DNMT3A*, with fitness values of 0.338 and 0.184 relative to a median fitness of 0.019 (Fig. 4A). *NPM1* and *DNMT3A* are well known driver mutations for AML [18]. Corroborating our finding, TreeMHN determined the *DNMT3A* mutation to cause the largest number of other mutations (8 pairs) while only inhibiting one mutation. In addition, CloMu determined mutations *ASXL1, GATA2* and *U2AF1* to have high fitness, with values ranging from 0.0394 to 0.0546. On the other hand, we found that *FLT3* had a below median fitness of 0.00585. This matches the fact that TreeMHN determined *FLT3* to have the largest number of strong negative causal relationships (3 pairs).

**Figure 4.**
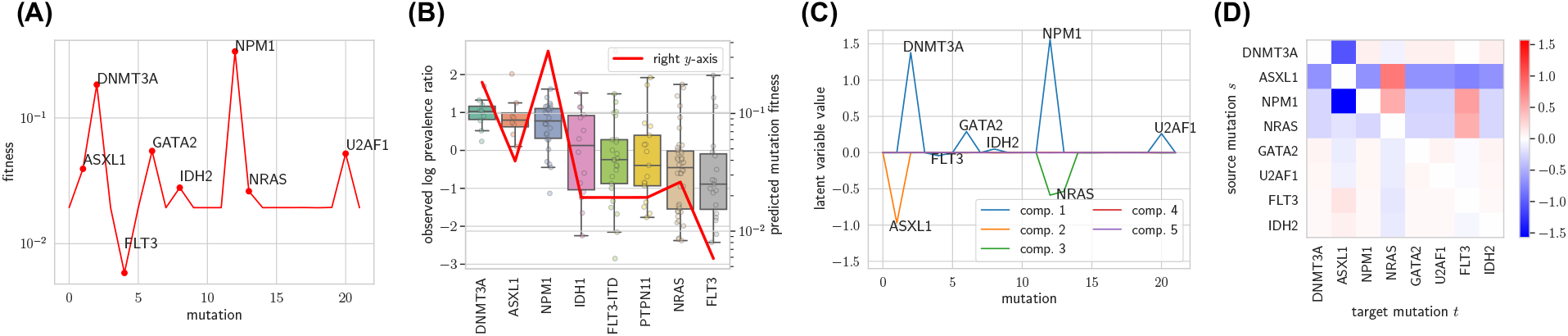
CloMu predicts fitness values validated by clone prevalence measurements and uncovers interchangeability and relative causal relationships on an AML dataset [11]. (A) CloMu identifies seven mutations which have a far greater fitness value than other mutations, and identified one mutation (*FLT3*) with a substantially lower fitness value than other mutations. (B) Box plot of log prevalence ratios for the eight mutations with the low standard error in log prevalence ratios (left *y*-axis), as well as a line showing our predicted mutation fitness values (right *y*-axis). The results demonstrate the validity of our fitness predictions. (C) Latent representations of mutations obtained from the *L* = 5 hidden neurons. This plot shows that *GATA2* and *U2AF1* are interchangeable on these data. (D) Relative causal relationships between mutation pairs (*s, t*). A value greater than 0 (red) is indicative of mutation *s* (row) causing mutation *t* (column) in the same clone, whereas a value smaller than 0 (blue) is indicative of mutation *s* inhibiting the occurrence of mutation *t* in the same clone.

To confirm the validity of our fitness predictions, we investigated the clone prevalence data that we did not use for training CloMu. Specifically, for each patient *p* and mutation *s*, let *γ*_1_(*p, s*) be the prevalence of the clone in which mutation *s* was introduced. Additionally, let *γ*_0_(*p, s*) be the prevalence of the parent clone of the clone in which mutation *s* was introduced. We define the *log prevalence ratio* as Γ(*p, s*) = log((*γ*_1_(*p, s*) + 0.01)*/*(*γ*_0_(*p, s*) + 0.01)), which adds 0.01 to all prevalence measurements in order to avoid issues caused by dividing by or taking the log of very low numbers. Intuitively, if mutation *s* is highly fit it should cause the clone that introduced mutation *s* to outgrow its parent clone, which would lead to a positive value for Γ(*p, s*). Conversely, a mutation *s* that does not increase fitness of the clone that introduced it would lead to a non-positive Γ(*p, s*). We analyzed the eight mutations with the lowest standard error in their Γ(*p, s*) measurements. Three of these mutations (*DNMT3A, ASXL1* and *NPM1*) were determined to be highly fit by our model and five were determined to have low fitness (*IDH1, FLT3-ITD, PTPN11, NRAS* and *FLT3*). The log prevalence ratio measurements agreed with our model’s conclusion on all eight mutations (Fig. 4B). In fact, the one mutation *FLT3* which our model predicted to have below median fitness also was determined to have the smallest median log prevalence ratio of −0.888.

CloMu also analyzed shared properties and interchangeability of mutations. We found *GATA2* and *U2AF1* to be interchangeable with similar latent representations (Fig. 4C). Additionally, we found *NPM1* and *DNMT3A* to have shared properties. Specifically, the first dimension of their latent representation is nearly identical (shown in blue in Fig. 4C). However, *NPM1* also has a shared property in the third latent dimension (showed in green in our plot), which is not present in *DNMT3A*. Finally, *ASXL1* was determined to be completely unique, utilizing the second dimension in its latent representation (showed in orange in our plot) unlike any other mutation. These patterns of interchangeability were also reflected in the causality relationships identified by CloMu (Fig. 4D).

The two main positive causal relationships found are from *ASXL1* and *NPM1* to *NRAS* — with *R*(*ASXL1, NRAS*) = 0.814 and *R*(*NPM1, NRAS*) = 0.505 (Fig. 4D). TreeMHN agrees with our conclusion that *NPM1* causes *NRAS*. Although GeneAccord detects the co-occurrence of *NPM1* and *NRAS*, the method does not have the capability to identify the directionality causal relationships. In our model, we found the strongest negative causal relationship to be from *NPM1* to *ASXL1*, i.e. *R*(*NPM1, ASXL1*) = −1.57. In addition, we identified a strong negative causal relationship in the reverse direction, i.e. *R*(*ASXL1, NPM1*) = −0.667. Such bidirectional negative causality is indicative of mutual exclusivity. Indeed, TreeMHN also identified this bidirectional negative causal relationship. Mutual exclusivity between *ASXL1* and *NPM1* has been described previously in the literature [20]. Moreover, a strong negative association from *ASXL1* to *FLT3-ITD* has also been reported [20]. Indeed, CloMu identified a strong negative causal relationship from *ASXL1* to *FLT3-ITD* (*R*(*ASXL1, FLT3-ITD*) = −0.754 and *R*(*FLT3-ITD, ASXL1*) = 0.131).

## 5 Discussion

In this work, we introduced CloMu, a tree-generative model of cancer evolution, which can be used to perform tree selection, determine mutation fitness, causality, interchangeability and detect complex evolutionary pathways composed of sets of interchangeable mutations. Like TreeMHN [1] and HINTRA [2], CloMu models the generation of trees by independently determining the rate of new mutations occurring on a clone based on the clone’s current mutations. CloMu uses a low-parameter neural network, which affords it the flexibility to model complex multi-mutation interactions while retaining resistance to overfitting. Using simulations without interchangeable mutations, we demonstrated that CloMu matches TreeMHN’s and outperforms all other baseline methods for the task of detecting causal relationships. Additionally in the presence of interchangeable mutations CloMu greatly outperformed TreeMHN as well. Our simulations further demonstrated CluMu’s accuracy in performing tree selection and identifying interchangeable mutations and their evolutionary pathways, outperforming competing methods. On real cancer data consisting of a breast cancer cohort [11] and an acute myeloid leukemia (AML) [12], CloMu was able to assess mutation fitness, determine interchangeable mutations and find causal relationships. We found that mutations with high fitness typically correspond to known driver mutations, and that causal relationships matched previously reported associations and patterns of mutual exclusivity. For the AML data, we performed additional orthogonal validation by comparing predicted mutation fitness values to prevalences of clones, finding high consistency. These finding demonstrate that CloMu is a versatile and effective method for a wide variety of prediction tasks regarding cancer evolution.

There are several directions for future work and expansions of CloMu. First, one could use CloMu’s lowdimensional representations of clones and mutations to predict measured properties of cancer clones or mutations, such as response to treatment or other patient outcomes. Second, one could extend CloMu’s model to support somatic mutations beyond single-nucleotide variants (SNVs) such as copy-number aberrations and their interplay with SNVs. Third, we only evaluated multi-mutation effects on simulated data, which required a large number *n* ≈ 500 of patients and large effect sizes. Our simulations showed that it is feasible to detect complex patterns such as two mutations *s* and *t* only causing a third mutation *r* when paired together. To perform such analyses on real data in future work, we require much larger data sizes than currently available, which we expect to see in the future.

## Acknowledgements

We thank the Beerenwinkel Lab and Xiang Ge Luo for providing access to TreeMHN simulation results. This work started as a course project in “CS598MEB: Computational Cancer Genomics”. M.E-K. was supported by the National Science Foundation (CCF-2046488) as well as funding from the Cancer Center at Illinois.

---

**Figure S1.**
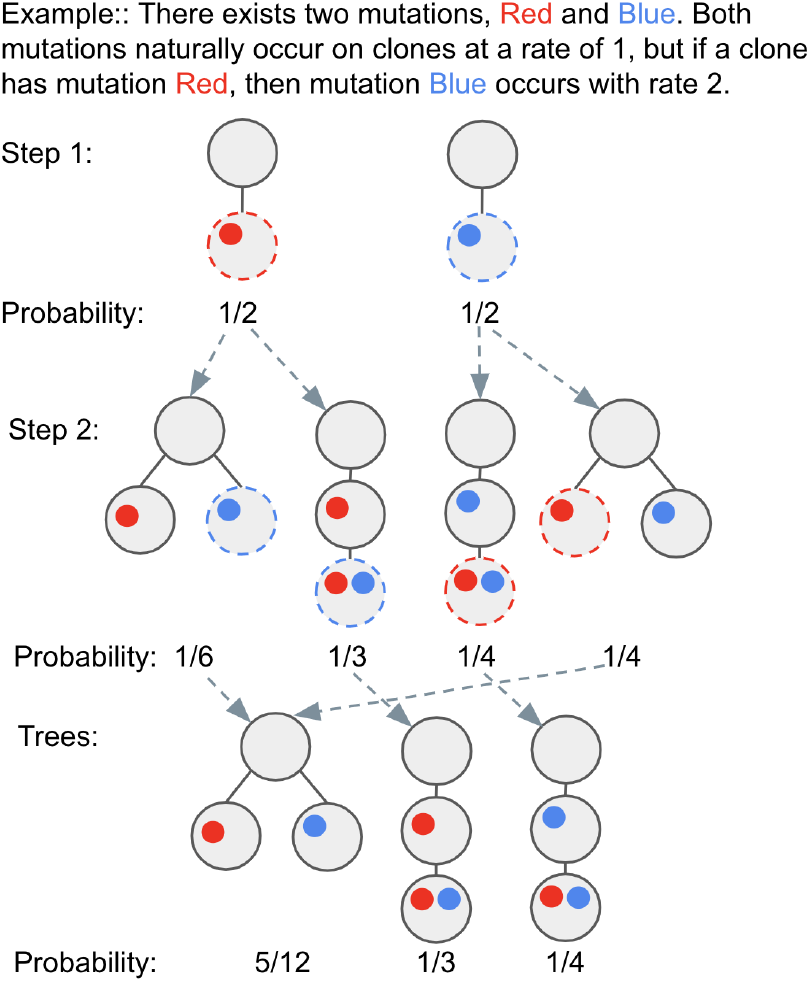
This is a toy example to demonstrate how causal relationships effect the probability of phylogeny trees in a simple system. It also demonstrates how there are multiple ways of generating the same tree; specifically, trees 1 and 4 in step 2 are the same tree.

## A Supplementary Methods

### A.1 CloMu: Neural Network

CloMu uses a two layer neural network with a small number *L* of hidden neurons for the function *f*_*θ*_. Specifically, the input to the neural network is the vector **c** and the output is a rate *f*_*θ*_(**c**, *s*) for each mutation *s*. This model has 2*Lm* + *m* + *L* = *O*(*Lm*) total parameters. We use a tanh nonlinearity rather than a ReLU nonlinearity to reduce the risk of latent variables being stuck at zero during training. To optionally enforce the infinite sites assumption, we set *f*_*θ*_(**c**, *s*) to a very large negative number if *c*_*s*_ = 1 to ensure a mutation is not added twice to the same clone.

To enable easier regularization, which is used primarily for the sake of interpretability but also used to avoid overfitting on some extremely small datasets, we made an two additional modifications to the neural network. First, the neural network also contains a single scalar value for each input mutation, which is added to all variables in the output equally. This allows one variable per mutation to influence the general fitness of the clone, allowing for a mutation to affect clonal fitness while only modifying one parameter (rather than modifying one parameter per output mutation). Specifically, this is equivalent to adding a function independent of *s, h*_*θ*_(**c**), to *f*_*θ*_(**c**, *s*) like the below equation.

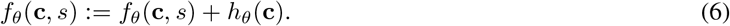

Second, a one-hot encoded variable representing the number of mutations in the clone is included. This allows the model to take into account the number of mutations on a clone without modifying the parameter for every individual input mutation. This therefore allows the model to easily know if the clone is the non-mutated root clone with only one parameter.

These two modifications are essentially useless and give no empirical advantage when no regularization is used, since these variables can automatically be computed and used from the already existing data. However, once L1 regularization is included during training, these variables help the model achieve sparsity. On very small datasets sparsity can be important to avoid overfitting; however, on larger datasets, sparsity is simply useful because it helps with interpretability. Having all variables that do not need to be nonzero set to zero helps with creating good plots and thus interpretability. This also likely helps with any downstream tasks that would use the latent representation of mutations or clones generated by the model.

### A.2 Determining Evolutionary Pathways

For the sake of interpretability, it is useful to represent patterns in cancer evolution as evolutionary pathways that frequently occur. We define an evolutionary pathway of interchangeable mutations as a list of sets of mutations such that the cancer obtained one mutation from each set in order. More precisely, define an *evolutionary pathway* of length, 𝓁 as sets of mutations *S*_1_, …, *S*_𝓁_. A clone with mutations *s*_1_, …, *s*_𝓁_ in temporal order fits the evolutionary pathway *S*_1_, …, *S*_𝓁_ if *s*_*k*_ ∈ *S*_*k*_ for all *k* ∈ [𝓁].

The probability of an evolutionary pathway *S*_1_, …, *S*_*𝓁*_ is defined as the probability that a clone with 𝓁 mutations fits the evolutionary pathway. Define **c**(*s*_1_, …, *s*_*k*_) as the clone with mutations *s*_1_, …, *s*_*k*_. The probability of the evolutionary pathway *S*_1_, …, *S*_*𝓁*_ is given by the below expression.

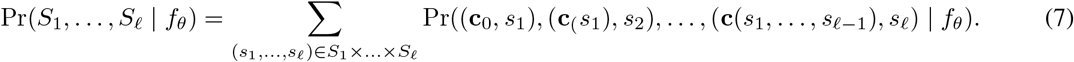

As a null model, we consider the probability Pr_0_(*s* | *f*_*θ*_, 𝓁) of a mutation *s* occurring in a pathway of length 𝓁.

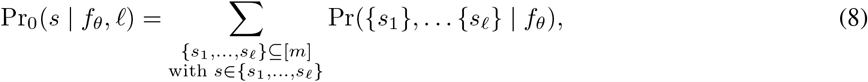

where Pr({*s*_1_ }, …, {*s*_𝓁_} | *f*_*θ*_) is the probability of the pathway {*s*_1_}, … {*s*_𝓁_}.

The null probability, ignoring causal relationship, of a mutation *s* occurring at a particular step for a particular clone would be Pr_0_(*s* | *f*_*θ*_, 𝓁)*/*𝓁. The null probability for a pathway would then be the null probability of generating every clone that fits the pathway based on the default probability in every step of generating the clone. Specifically, the null probability of a pathway *S*_1_, …, *S*_𝓁_ is defined by the below equation.

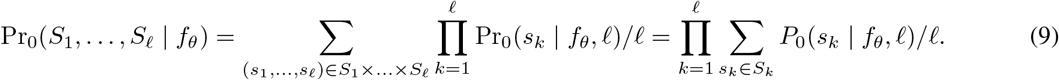

The expression Pr(*S*_1_, …, *S*_𝓁_ | *f*_*θ*_)*/* Pr_0_(*S*_1_, … *S*_𝓁_ | *f*_*θ*_) defines how much more likely a pathway is to to occur than what one would expect if there were no causal relationships. For a pathway to be meaningful, one wants Pr(*S*_1_, …, *S*_𝓁_ |*f*_*θ*_)*/* Pr_0_(*S*_1_, …, *S*_𝓁_ |*f*_*θ*_) to be high in order to indicate that the pathway demonstrates meaningful causal relationships. Additionally, one wants Pr(*S*_1_, …, *S*_𝓁_ | *f*_*θ*_) itself to be high so that the pathway occurs frequently enough to be meaningful. Define the score of a pathway with the below equation, where *a >* 0 is a nonnegative constant.

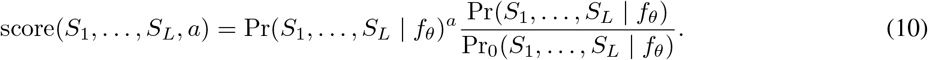

The value of *a* can be adjusted based on the specific application. When implementing this we operate in logarithmic space.

In order to determine the most meaningful pathways, one can optimize for the score of the pathway. Specifically, given a length 𝓁 ≤ *m* the procedure used in this paper starts with an evolutionary pathway *S*_1_, …, *S*_𝓁_ where each *S*_*k*_ = [*m*]. Then, in a greedy fashion, we add or remove a single mutations that maximizes score(*S*_1_, …, *S*_𝓁_, *a*). We repeat this until a locally optimal pathway is found. Once an individual optimal pathway is found, we set the probability Pr((**c**_0_, *s*_1_), …, (**c**(*s*_1_, …, *s*_𝓁−1_), *s*_*𝓁*_) | *f*_*θ*_) used in (7) to 0 in a lookup table so as to discourage repeatedly finding the same or similar pathways. Then, the optimization procedure is run again to generate a new pathway. If the score of a pathway reaches below Pr([*m*], …, [*m*] | *f*_*θ*_)^*a*^, it is not at all meaningful and the process of generating optimal pathways ends. This cut off is used since the pathway consisting of only the set of all mutations at every step has that score.

### A.3 CloMu: Reinforcement Learning

In this section we discuss how we used reinforcement learning to identify model parameters *f*_*θ*_ for the the neural network. Specifically, Appendix A.3.1 gives a brief introduction to policy learning. In Appendix A.3.2 we discuss how policy learning can be applied to CloMu’s model. Finally, in Appendix A.3.3 we discuss a speedup for the typical case of a small number of trees per patient.

#### A.3.1 Basic Introduction to Policy Learning

The type of reinforcement learning used in this paper is policy learning. Define *A* as a sequence of actions the model may take, *R*(*A*) as the reward for those actions, and Pr(*A* | *f*_*θ*_) as the probability our model takes those actions. In our application, *A* will be the phylogenetic generation processes sampled from our model. Define *S*_*A*_ as the set of all possible sequences of actions our model could take. In our application, *S*_*A*_ would correspond to the set of all tree generating processes. To optimize the reward that the model gets, one wants to increase the probability of high reward actions. Therefore, one wants to maximize the below equation.

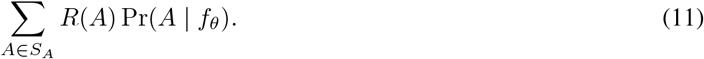

The ideal gradient for optimizing this would theoretically be the below equation.

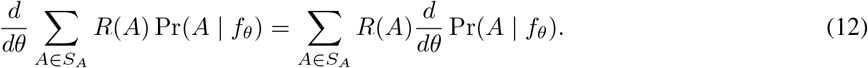

In practice it is infeasible to sum over all *A* ∈ *S*_*A*_ since *S*_*A*_ may be a very large set. However, one can estimate (12) by sampling from Pr(*A* | *f*_*θ*_):

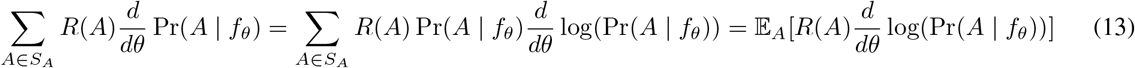

In the typical case where *R*(*A*) is independent of *θ* this gives the below reward function to optimize.

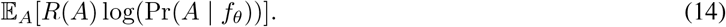

#### A.3.2 Policy Learning for CloMu

In our application, a sequence *A* of actions will be the phylogenetic generation processes *G* sampled from our model. Moreover, the set *S*_*G*_ of all possible sequences of actions corresponds to the set of all tree generating processes on *m* mutations. To use reinforcement learning, we need to meet two criteria. First, we need to be able to sample from our distribution Pr(*G* | *f*_*θ*_). For our model, this is very easy since *G* is generated by a simple probabilistic process (Section 2). Second, we need to be able to formulate our objective as the below equation, where *R*(*G*) is the reward function.

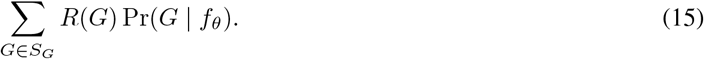

An important aspect of *R*(*G*) is that the gradient with respect to *θ* that is used for optimization does not affect *R*(*G*). To make this explicit and clear, instead of formulating our objective function as that objective function, we formulate the gradient of our objective function to fit the reinforcement learning objective function gradient. Specifically, to apply policy learning, the gradient of our objective function must be formulated as below.

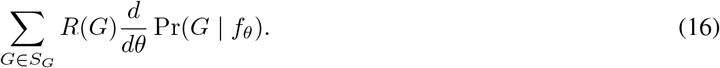

We can do so through the below manipulations of the gradient of the log likelihood.

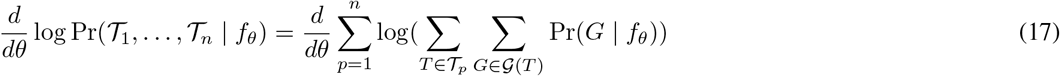

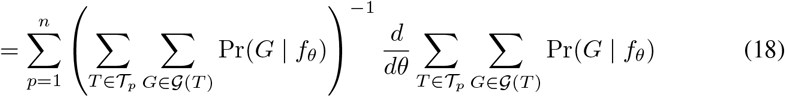

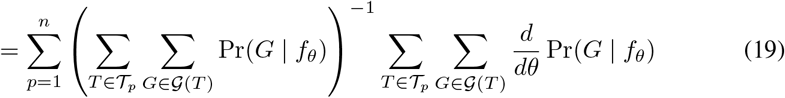

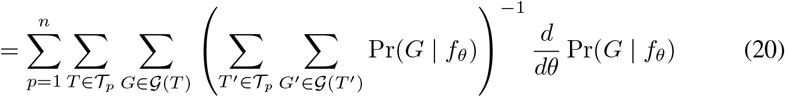

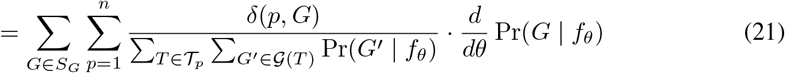

In the above derivation, we define *δ*(*p, G*) as 1 if *G* ∈ 𝒢 (*T*) for some *T* ∈ 𝒯_*p*_, and *δ*(*p, G*) = 0 otherwise. In other words *δ*(*p, G*) indicates whether *G* corresponds to an observed tree of patient *p*. Equation (21) corresponds to the reinforcement learning gradient if we set the reward function as below.

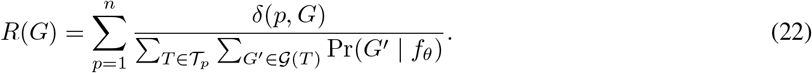

In reinforcement learning, one is sampling from Pr(*G* | *f*_*θ*_), so we can easily estimate ∑_*T*_ ∈𝒯_*p*_∑_*G*_∈𝒢 (*T*) Pr(*G* | *f*_*θ*_) by simply observing what fraction of generated *G* are in 𝒢 (*T*) with *T* ∈ 𝒯_*p*_. In other words, we can observe what fraction of generated trees are possible for patient *p*. This fraction changes as we train the model, and therefore must constantly be updated. However, assuming the model does not change much in a small number of iterations, it does not need to be calculated from scratch in every gradient update. The reinforcement learning objective function is now the below equation where *R*(*G*) is treated as a numerical value and no gradient is applied to it. In PyTorch this can be done by applying `.detach()`to *R*(*G*).

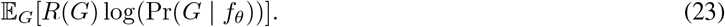

#### A.3.3 Policy Learning with Few Trees Per Patient

When there are very few trees per patient, it can be useful to directly model the probability of each individual tree, rather than indirectly modeling the probability of each tree through reinforcement learning. In particular, for such data, the reinforcement learning method introduced in the previous section may struggle to find those trees, and would perform much better if the probability of each tree was explicitly modeled. However, one cannot just avoid using reinforcement learning in those cases because there could still be a very large number of generation processes per tree. To solve both of these problems, we use a modified reinforcement learning system where the probability of each possible tree is explicitly modeled, but the probability of each generation process is not explicitly modeled and only learned through reinforcement learning.

To do so, for each possible tree *T* in the dataset (the union of 𝒯_*p*_ for all patients *p* ∈ [*n*]), we only sample generation processes *G* ∈ 𝒢(*T*). Of course, this requires a modification of the objective function to compensate for the change in the sampling procedure. Define *P*_*T*_ (*G* | *f*_*θ*_) as the probability of generating *G* with our procedure restricted such that it always generates processes which give the tree *T*. Define *R*′(*G*) = *R*(*G*) Pr(*G* | *f*_*θ*_)*/P*_*T*_ (*G* |*f*_*θ*_). *R*′(*G*) is the new reward function for this adjusted sampling procedure as shown in the below calculation.

**Table S1:**
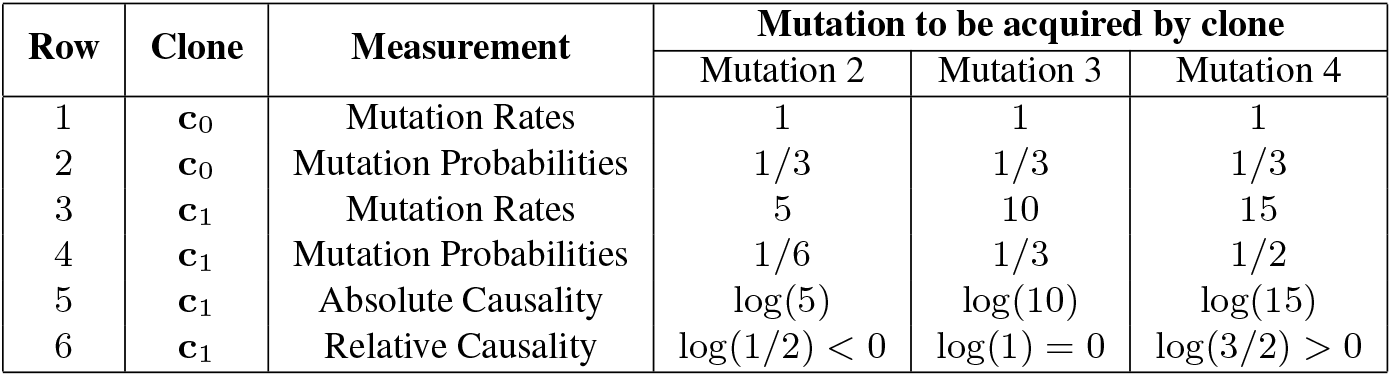
This demonstrates absolute causality and relative causality in a theoretical dataset with 4 mutations. Specifically, it shows the absolute and relative causality of a hypothetical mutation 1 measured using its clone **c**_1_. For the sake of simplicity, we assume mutation 1 occurs with a rate of 0 on both clones. Row 2 is calculated from row 1 by dividing each column by the sum of the three columns. The same calculation gives row 4 from row 3. Row 5 is the log of row 3 divided by row 1. Row 6 is the low of row 4 divided by row 2. Since mutation 1 is highly fit, it causes all other mutations in terms of absolute causality. However, in terms of relative causality, it causes mutation 4, inhibits mutation 2 and has no effect on mutation3.

**Figure S2.**
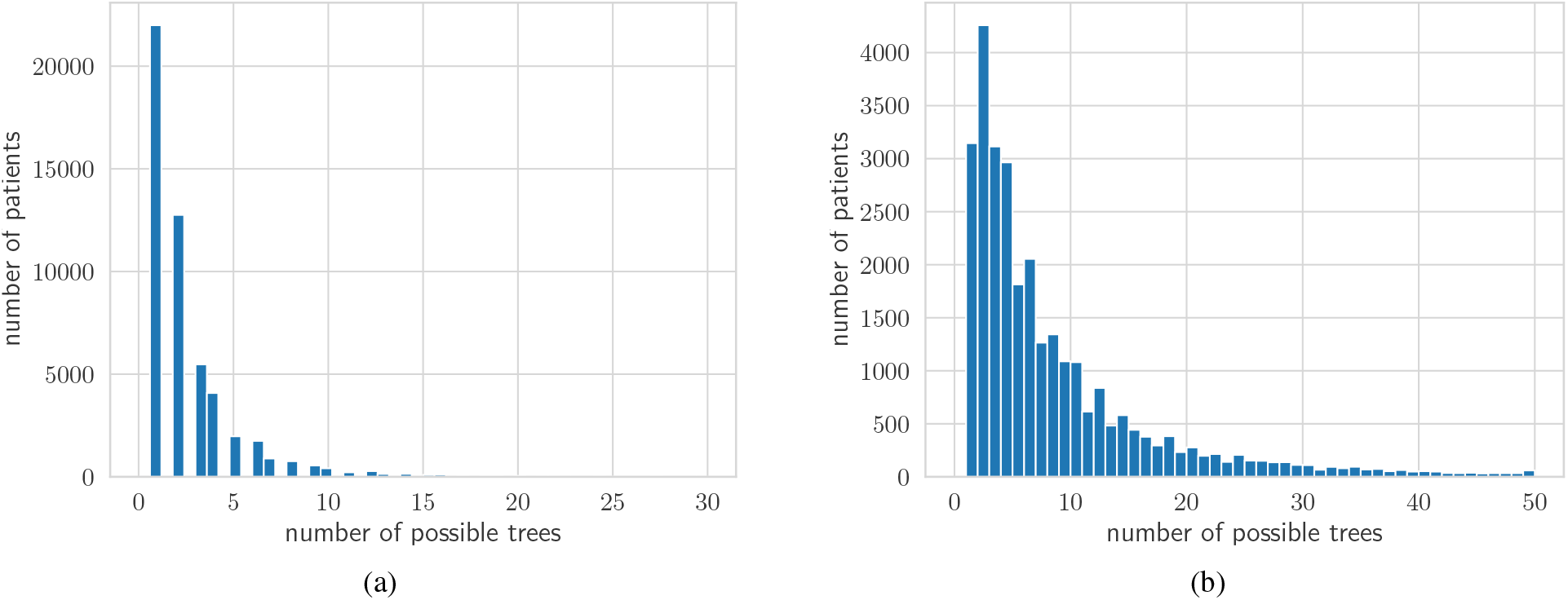
Number of trees in simulated data. Histogram of the number of trees for each patient in (a) Simulations (i) used to assess tree selection (b) Simulations (ii) used to assess mutation interchangeability and evolutionary pathway identification.

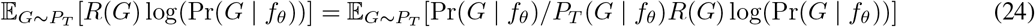

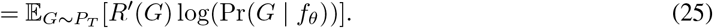

## B Supplementary Results

### B.1 Simulations

#### B.1.1 Setup for Simulations (I-b), (I-c) and (II)

To simulate causality with interchangeable mutations, we consider each mutation as a driver mutation. We partition the set of *m* mutations into 5 pairwise disjoint sets *S*_1_, …, *S*_5_ of equal size. For each pair (*S*_*i*_, *S*_*j*_) of mutation sets (where *i, j* ∈ {1, …, 5}), we determined *S*_*i*_ as causing *S*_*j*_, *S*_*i*_ as inhibiting *S*_*j*_ and *S*_*i*_ and *S*_*j*_ having no causal effect with uniform probabilities of 1*/*3 for each case. With a slight abuse of notation, we set the rate multiplier *g*(*S*_*i*_, *S*_*j*_) to 11 if *S*_*i*_ causes *S*_*j*_, 1*/*11 if *S*_*i*_ inhibits *S*_*j*_, and 1 if there is no causal relationship from *S*_*i*_ to *S*_*j*_.. The resulting rate multiplier *g*(*s, t*) of mutations *s, t* ∈ [*m*] equals the rate multiplier *g*(*S*_*i*_, *S*_*j*_) of their comprising sets (i.e., *s* ∈*S*_*i*_ and *t* ∈ *S*_*j*_). Similarly, we compute rates *f* (**c**, *t*) as in the Main Text and generate a ground-truth 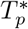 for each patient *p* ∈ [*n*] by first drawing the number 𝓁 of mutations of tree 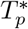 uniformly from {5, …, 7}. Subsequent bulk sequencing simulation with five sequencing samples resulted in sets 𝒯_1_, …, 𝒯_*n*_ of trees per patient (Fig. S2A). We varied the number *m* ∈ {10, 15} of mutations. Consequently, the size of each set *S*_*i*_ will either be 2 (for *m* = 10) or 3 (for *m* = 15). We simulated *n* = 400 patients for *m* = 10 and *n* = 600 patients for *m* = 20. For each setting of *m*, we generated 20 simulation instances. These instances are denoted as Simulations (I-b) and (I-c) (see Table 2).

**Table S2.** This table demonstrates how true positives, true negatives, false positives and false negatives are determined based on predicted and true causal relationships in the case of signed and bidirectional relationships.

| Predicted Relationship | True Relationship |  |  |
| --- | --- | --- | --- |
|  | No Relationship | Positive Relationship | Negative Relationship |
| No relationship | True Negative | False Negative | False Negative |
| Positive Relationship | False Positive | True Positive | False Positive |
| Negative Relationship | False Positive | False Positive | True Positive |

For evolutionary pathway assessment, we generated simulation instances that consists of one pathway (25% probability) or two pathways (75% probability). We considered *m* = 20 mutations, some of which will occur in pathways and are drivers and the remaining mutations are passengers. Each pathway consists of three pairwise disjoint sets *S*_1_, *S*_2_, *S*_3_ of mutations. In the case of two pathways all mutation sets are pairwise disjoint. The number of mutations in a set *S*_*i*_ is drawn uniformly from {1, 2, 3}. As such, the number of drivers ranges from 3 mutations (in the case of a single pathway with one mutation per set) to 18 mutations (in the case of two pathways with three mutations per set). To generate trees according to these evolutionary pathways we defined rates *f* (**c**, *t*) according to the stage and the pathway a particular clone is at. Specifically, clone **c** is at stage *i* ∈ { 0, …, 3 } of the pathway *S*_1_, *S*_2_, *S*_3_ if clone **c** contains at least one mutation from each set *S*_1_, …, *S*_*i*_ and does not contain a mutation from set *S*_*i*+1_ (here if *i* = 3 then *S*_4_ = ∅). We set the rate *f* (**c**, *s*) to 5 if **c** is at stage 0 for both pathways and *s* ∈*S*_1_ for some pathway. We set the rate *f* (**c**, *s*) to 20 if **c** is at stage *i >* 0 for some pathway and *s* ∈*S*_*i*+1_ for that pathway. If neither condition is met, we set rate *f* (**c**, *s*) = 1. Note, by definition, the normal clone **c** is at stage 0 for both pathways. Additionally, if a clone is at stage 3 then all mutations have rate 1. Similarly to the other simulations, we use these rates to generate a ground-truth 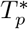 for each patient *p* ∈ [*n*] by first drawing the number 𝓁 of mutations of tree 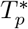 uniformly from {5, …, 7}. Subsequent bulk sequencing simulation with three sequencing samples resulted in sets 𝒯_1_, …, 𝒯_*n*_ of trees per patient (Fig. S2(b)). We set *n* = 500 patients, and generated a total of 30 simulation instances. These instances are denoted as Simulations (ii) (see Table 2).

#### B.1.2 Evaluating Bidirectional and Signed Causal Relationships

To evaluate bidirectional causality for signed causal relationships, one must calculate precision and recall from predicted and true causal relationships. First, every ordered pair of mutations is considered. Then, if some causal relationship (positive or negative) from *s* to *t* is predicted correctly, this is considered a true positive. Similarly, if no causal relationship is predicted from *s* to *t* and there is no true causal relationship from *s* to *t*, then this is considered a true negative. If there is a predicted causal relationship (positive or negative) from *s* to *t*, and this predicted relationship is incorrect for any reason, then it is considered a false positive. Note, this implies that if the predicted causal relationship is positive (or negative) and the true causal relationship is negative (or positive), then it is considered a false positive, despite their being a true causal relationship. This information is summarized in table S2.

Using this, each ordered pair of mutations can be classified as either a true positive, true negative, false positive or false negative. Summing these numbers across all ordered pairs of mutations gives total true positives, true negative, false positives and false negatives. Then, the precision is defined as the true positives divided by the sum of true positives and false positives. The recall is defined as the true positives divided by the sum of true positives and false negatives. These definitions follow those used in TreeMHN [1].

#### B.1.3 RECAP and TreeMHN Simulation Data

We benchmarked CloMu on previously published simulation datasets generated in the RECAP and TreeMHN papers. Specifically, for RECAP, we restricted the comparison to instances with (*m*, 𝓁) ∈ { (5, 5), (7, 12) } where *m* is the total number of mutations and 𝓁 is the number of mutations per patient. For TreeMHN we restricted to instances with *n* = 300 patients, and randomly selected 20 instances for each number *m* ∈ {10, 15, 20} of mutations.

### B.2 Method Parameters

- On Simulations (IV), TreeMHN was ran with default parameters and the use of stability selection. For comparison on Simulations (I), we ran TreeMHN with the penalty parameter set to maximize precision and recall. Additionally, we did not use stability selection, since improving precision at the cost of recall would not improve their results.
- We ran RECAP with the default parameters. For Simulation (II) we implemented a custom method of determining pathways of interchangeable mutations from RECAP.
- We ran REVOLVER with default parameters. For Simulation (II) we used the same custom implemented method as was used for RECAP.
- We ran GeneAccord with default parameters.
- We did not run HINTRA due to its lack of scalability. However, we did compare with HINTRA on RECAP’s data (Simulation (III)) which was ran with default parameters.

### B.3 Baseline Heuristics

For Simulations (I-a) and (II), we included RECAP and REVOLVER despite their implementation not directly supporting these tasks. RECAP and REVOLVER are consensus tree method (Main Text Table 1), which select one tree per patient.

To support causality inference for RECAP and REVOLVER, we applied a post-processing step. Specifically, we processed RECAP and REVOLVER’s selected trees for each patient from the set of possible trees. We then extracted all edges from these trees and determined the frequency of each edge. For Simulations (I-a), we considered there to be a positive causal relationship from mutation *s* to mutation *t* if the edge (*s, t*) occurred with frequency above a fixed threshold. We used a threshold of 40 based on the number *n* = 500 patients for Simulations (I-a).

For Simulations (II), the goal is to predict the edges which occur in the simulated pathways. Specifically, if a ground-truth pathway contains sets *S*_*i*_ and *S*_*i*+1_, we say that pathway contains all edges (*s, t*) with *s* ∈*S*_*i*_ and *t* ∈*S*_*i*+1_. Given a threshold frequency, RECAP/REVOLVER predict an edge to exist for a pathway in a simulation if that edge occurs in the selected trees with a frequency above the threshold. We used a threshold of 20 based on the number *n* = 500 patients for Simulations (II).

### B.4 Results on RECAP Simulations – Simulations (III)

The RECAP simulated dataset (denoted as Simulations III in Table 2) has a small number of patient subtypes each of which has a single phylogeny tree. Within the RECAP simulated data, two groups of simulated datasets are used. One group has 5 mutations per patient and 5 mutations total. The second group has 7 mutations per patient and 12 mutations total. For each dataset (in each group), there are *k* ∈ {1, …, 5} clusters among *n* ∈{50, 100} patients. Each patient cluster correspond to one true tree. In the published RECAP simulations, each patient was randomly assigned to a true tree, then the set of possible trees is generated by simulated bulk frequency measurements. The goal in this simulated dataset is to assign the correct tree to each patient.

#### B.4.1 Post-processing to Support Identification of Consensus Trees

Performing well on the RECAP dataset requires the knowledge that there are a small number of true trees in each dataset, which is not an intrinsic property of our model. To adjust for this fact, a post processing step to our model must be added. Specifically, tree predictions are generated for each patient according to the following procedure. As previously defined, let Pr(*T* | *f*_*θ*_) be the probability that tree *T* is generated according to the model, and 𝒯_*p*_ be the set of possible trees for patient *p*. For a set 𝒮 of trees define Pr(𝒮 | *f*_*θ*_) as the below equation.

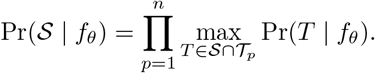

In other words, Pr(S | *f*_*θ*_) is the probability of the maximum likelihood possible trees (*T*_1_, …, *T*_*n*_) ∈ 𝒯_1_ × … ×𝒯_*n*_ when we restrict the set of trees to S. Thus, Pr(𝒮 | *f*_*θ*_) represents how well S works as the set of true trees.

Define 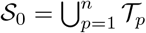 as the set of all input trees. Define 𝒮_*t*+1_ given 𝒮_*t*_ as follows.

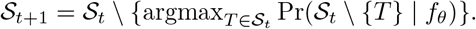

In other words, 𝒮_*t*+1_ is 𝒮_*t*_ with the element removed which decreases Pr(*f*_*θ*_) by the smallest quantity. Define *t*^*^ as the last *t* with 𝒯_*p*_ ∩ 𝒮_*t*_ non-empty for all *p*. Our final predicted tree 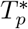 for each patient *p* is given by the below equation.

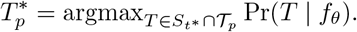

In simple terms, this procedure works by iteratively removing trees from the set of possible trees while predicting the maximum likelihood tree for each patient until no more trees can be removed from the set of possible trees. This then results in predictions 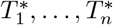.

#### B.4.2 Results

We compared CloMu to RECAP as well as REVOLVER. As in the Main Text, we consider the tree selection accuracy, which is the fraction of patients for which the correct true tree is predicted. Fig. S3 shows that CloMu performed almost identically to RECAP and far outperforms REVOLVER. In fact, our method and RECAP both achieved a 98.6% accuracy on average for simulations with 5 mutations and an average 100% accuracy for 12 mutations (Fig. S3A,C). We additionally investigated the predicted number of patient clusters for our method and RECAP, finding perfect performance for CloMu and near perfect performance for RECAP across varying number of mutations (Fig. S3B,D).

### B.5 Results on TreeMHN Simulations – Simulations (IV)

CloMu is generally competitive with but slightly less effective than TreeMHN on datasets from the TreeMHN paper. Datasets in the TreeMHN paper use completely independent causal relationships between all mutations, with no related or interchangeable mutations. In this case, using CloMu with a neural network function creates a slight disadvantage on datasets with a very poor signal-to-noise ratio. Therefore, we included a version of CloMu where the neural network function is replaced with a simple linear model, denoted as “Linear CloMu”. One virtue of the training setup of CloMu is that it is completely independent of the function chosen for *f*_*θ*_. Therefore, this only requires a minor tweak to the code. We compared CloMu to TreeMHN with stability selection in the cases of *n* = 300 patients, and *m* ∈ {10, 15, 20} mutations. The TreeMHN setup and hyperparameters were set exactly as in the TreeMHN paper. Like in the TreeMHN paper, we plot the precision and recall curves. Unlike the TreeMHN paper, we report the precision and recall on all mutations, not only the top half most frequent mutations.

Fig. S4 shows the precision and recall curves averaged over the 20 simulations per simulation setup. One can see that TreeMHN outperformed CloMu on these data, but CloMu remained competitive especially when a linear function is used for *f*_*θ*_. For instance, it seems CloMu with a linear model may extend to higher recall values more effectively than TreeMHN, but this is unclear since the range of hyperparameters used in the TreeMHN paper do not extend into high recall. From these results we conclude the following. If one knows for certain that there are no shared properties between mutations and that all causal relationships are independent, then TreeMHN is a slightly more effective method than CloMu. However, if there exists shared properties between mutations, CloMu ranges from slightly more effective to vastly more effective than TreeMHN.

**Figure S3.**
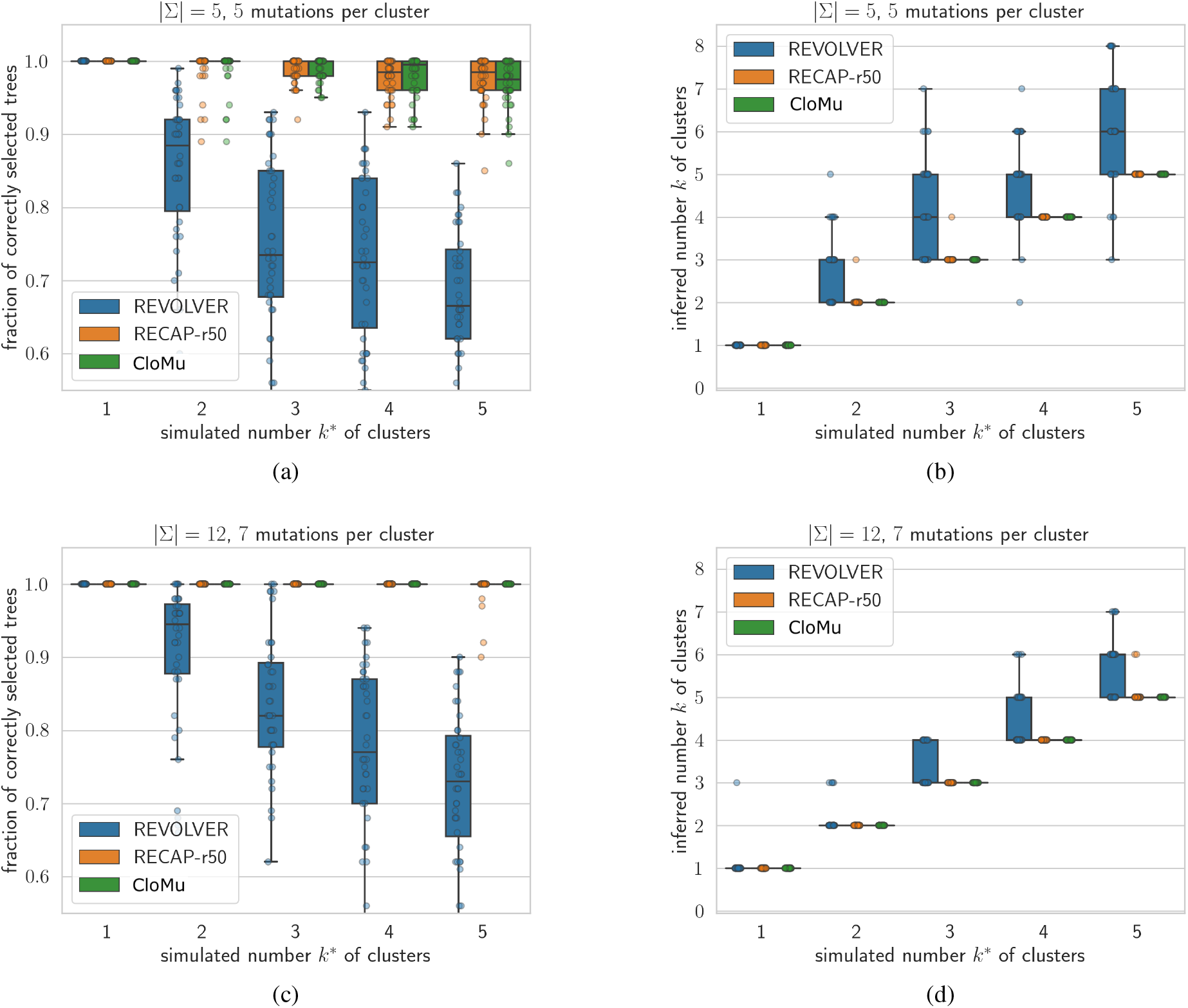
CloMu’s matches RECAP’s performance on RECAP’s simulated data (Simulations (III)). These plots show RECAP results with 50 restarts and additionally includes REVOLVER results reported in [3]. (A,C) Tree selection accuracy. (B,D) The number of ground-truth patient clusters vs. the predicted number of patient clusters. Panels (A-B) show results for |Σ| = 5 total mutations and *m* = 5 mutations per patient whereas panels (C-D) show results for |Σ| = 12 total mutations and *m* = 5 mutations per patient.

**Figure S4.**
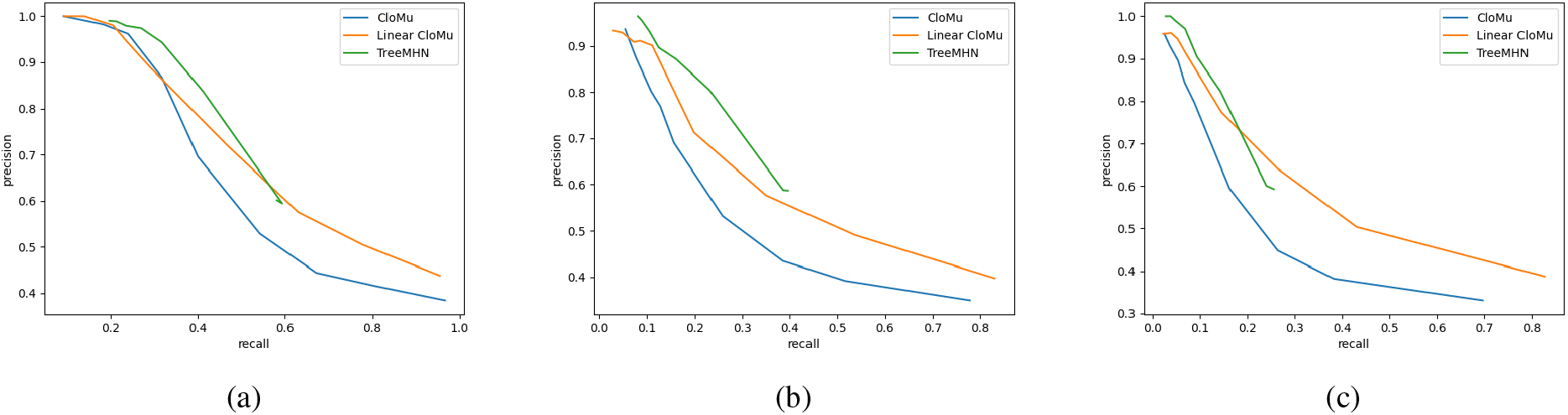
CloMu performs slightly worse than TreeMHN on TreeMHN’s simulated data (Simulations (IV). (A) Simulations with *m* = 10 total mutations. (B) Simulations with *m* = 15 mutations. (C) Simulations with *m* = 20 total mutations.

**Figure S5.**
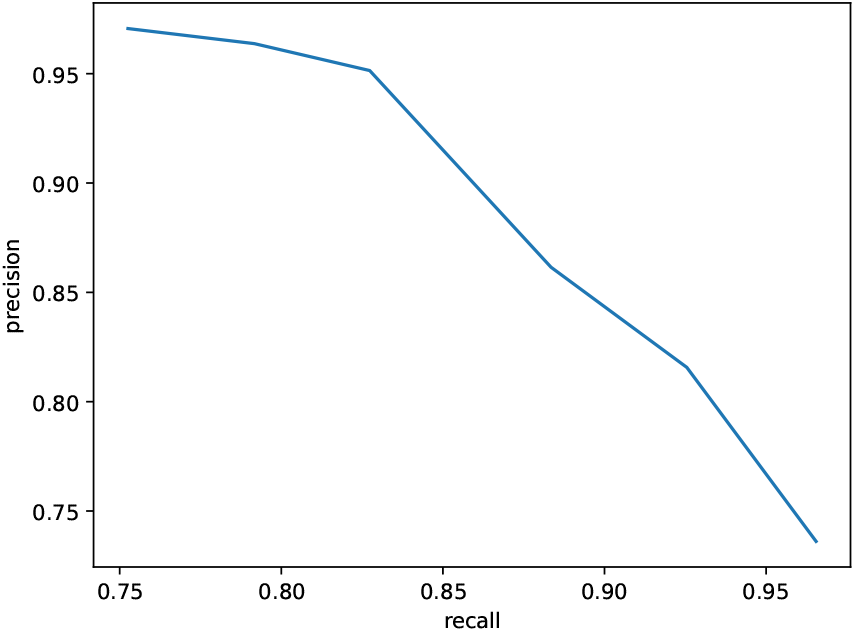
CloMu is robust to violations of the independent clonal evolution assumption. We consider a modification of Simulations (I-a) that results in a violation of the assumption. The resulting precision and recall values are slightly reduced but remain very high, as for the original Simulations (I-a) results shown in Main Text Fig. 2B).

### B.6 CloMu is Robust to Violations of the Independent Clonal Evolution Assumption

CloMu is built off of the independent clonal evolution assumption that only the mutations on a clone will effect the probability of new mutations occurring on that clone. This assumption may be generally realistic, but could have exceptions in real tumor evolution. Therefore, ideally we would want CloMu to be somewhat robust to this possibility. We test this by using a simulation where the causal relationships between mutations also occur to a smaller extent between mutations in different clones. In other words, if mutation *s* being in a clone causes mutation *t* to be much more likely to occur in that clone, then mutation *s* being in any clone will cause mutation *t* to be somewhat more likely to occur in any, not necessarily the same, clone.

We find that CloMu still performs well on a causal relationship simulation when clonal interactions are included in the simulation. Specifically, our 20 simulation instances result from a modified version of Simulations (I-a) with no interchangeable mutations, 5 driver mutations and 5 passenger mutations. If mutation *s* causes mutation *t*, then as a higher proportion of clones have mutation *s*, the probability of mutation *t* on all clones will increase. Specifically, if a proportion *p* of clones have some mutation *s*, then the log rate of mutation *t* will be increased by *p* log(2). Applying CloMu on this simulation gives causality precision-recall curve shown in Fig. S5, with values close to 1, similar to the original Simulations (I-a) results shown in Main Text Fig. 2B). Hence, for this scenario CloMu was robust to violations of the independent clonal evolution.

